# Identification and characterization of zebrafish Tlr4 co-receptor Md-2

**DOI:** 10.1101/817528

**Authors:** Andrea N. Loes, Melissa N. Hinman, Dylan R. Farnsworth, Adam C. Miller, Karen Guillemin, Michael J. Harms

**Affiliations:** Institute of Molecular Biology, University of Oregon, Eugene, OR 97403; Institute of Neuroscience, University of Oregon, Eugene, OR 97403; Department of Chemistry and Biochemistry, University of Oregon, Eugene, OR 97403; Department of Biology, University of Oregon, Eugene, OR 97403; Humans and the Microbiome Program, CIFAR, Toronto, Ontario M5G 1Z8, Canada

## Abstract

The zebrafish (*Danio rerio*) is a powerful model organism for studies of the innate immune system. One apparent difference between human and zebrafish innate immunity is the cellular machinery for LPS-sensing. In amniotes, the protein complex formed by Toll-like receptor 4 and myeloid differentiation factor 2 (Tlr4/Md-2) recognizes the bacterial molecule lipopolysaccharide (LPS) and triggers an inflammatory response. It is believed that zebrafish have neither Md-2 nor Tlr4: Md-2 has not been identified outside of amniotes, while the zebrafish *tlr4* genes appear to be paralogs, not orthologs, of amniote *TLR4s*. We revisited these conclusions. We identified a zebrafish gene encoding Md-2, *ly96*. Using single-cell RNA-Seq, we found that *ly96* is transcribed in cells that also transcribe genes diagnostic for innate immune cells, including the zebrafish *tlr4*-like genes. Unlike amniote *LY96*, zebrafish *ly96* expression is restricted to a small number of macrophage-like cells. In a functional assay, zebrafish Md-2 and Tlr4a form a complex that activates NF-κB signaling in response to LPS, but *ly96* loss-of-function mutations gave little protection against LPS-toxicity in larval zebrafish. Finally, by analyzing the genomic context of *tlr4* genes in eleven jawed vertebrates, we found that *tlr4* arose prior to the divergence of teleosts and tetrapods. Thus, an LPS-sensitive Tlr4/Md-2 complex is likely an ancestral feature shared by mammals and zebrafish, rather than a *de novo* invention on the tetrapod lineage. We hypothesize that zebrafish retain an ancestral, low-sensitivity Tlr4/Md-2 complex that confers LPS-responsiveness to a specific subset of innate immune cells.

## INTRODUCTION

Amniote innate immune systems are exquisitely sensitive to lipopolysaccharide (LPS), a component of the cell wall in Gram-negative bacteria (1–3). LPS is sensed by a protein complex composed of Toll-like receptor 4 (Tlr4) and Md-2 (also known as LY96 and ESOP-1) (1, 4). LPS binds in a pocket of Md-2, triggering dimerization of the Tlr4/Md-2 complex (Fig 1) (5). This, in turn, activates a Myd88-dependent NF-κB response (6). When properly regulated, the LPS activation of Tlr4/Md-2 regulates microbiome populations (7), recruits neutrophils to sites of infection (8), and induces angiogenesis (9). When dysregulated, Tlr4/Md-2 activity induces septic shock (10, 11), plays roles in inflammatory disorders (11, 12), and is a key player in the tissue remodeling that accompanies tumorigenesis (13, 14).

**Fig 1.**
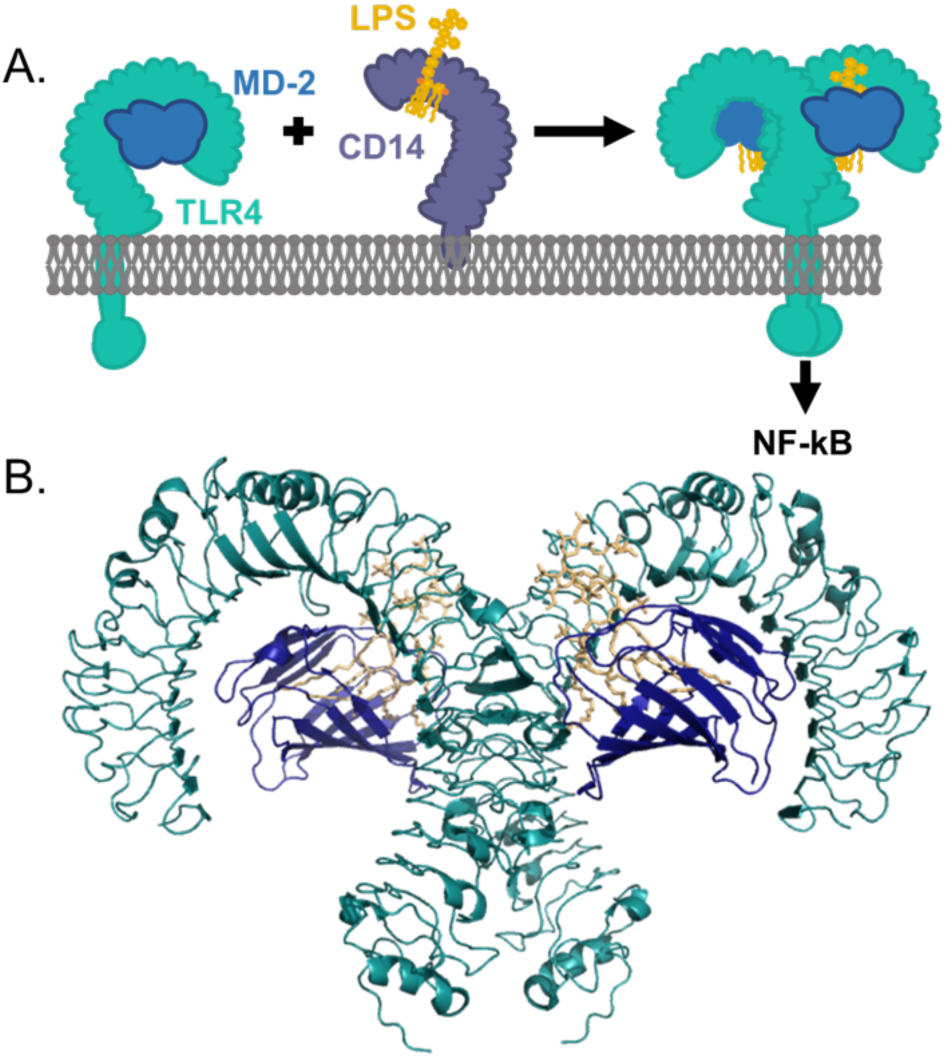
LPS activation of amniote Tlr4 requires cofactors Md-2 and Cd14. A) Schematic representation of LPS transfer from Cd14 to the Tlr4/Md-2 complex. LPS (yellow) is brought by Cd14 (purple) and loaded into Md-2 (navy blue). Md-2 is bound by Tlr4 (cyan). Binding of LPS to the Md-2 co-receptor causes dimerization of the Tlr4/Md-2 complex, activating a downstream inflammatory response. B) The interface between human Tlr4 (cyan) and Md-2 (navy blue) is extensive. Both are required to form a productive interaction with LPS (yellow). Structure shown was made from PDB 3FXI (19).

The role of Tlr4/Md-2 in LPS-sensing outside of amniotes remains poorly understood. Understanding this response in zebrafish (*Danio rerio*) is of particular interest, as the zebrafish is a powerful model organism for studies of vertebrate innate immunity (15). Zebrafish have mature genetic resources, rapid generation time, clear embryos, and facile germ-free derivation (16, 17). The zebrafish is increasingly popular as a model for understanding host-microbe interactions (18), as well as a tool to understand the development of the innate immune system (16).

The zebrafish response to LPS is puzzling (20, 21). In some ways it is similar to amniotes. As in amniotes, LPS triggers Myd88-dependent NF-κB inflammatory response (22, 23). Further, the expression patterns of genes induced by LPS stimulation are highly similar between mouse and zebrafish (24). There are, however, several lines of evidence that suggest Tlr4/Md-2 is not involved. Most critically, the gene encoding the essential co-receptor Md-2 has not been identified in zebrafish and other ray-finned fishes (20, 21, 25). Further, zebrafish Tlr4 proteins do not activate NF-κB in response to LPS in *ex vivo* assays, even when complemented with a mouse or human Md-2 (20, 21). Finally, zebrafish do not have a direct ortholog to amniote *tlr4*. Rather, they possess three *tlr4*-like genes—*tlr4ba*, *tlr4bb*, and *tlr4a1*—that are thought to have arisen from an ancestral Toll-like receptor lost in tetrapods but retained in ray-finned fishes (21). These observations have led to the hypothesis that zebrafish respond to LPS by a non-Tlr4/Md-2-dependent pathway.

We set out to carefully revisit these conclusions using resources unavailable when the initial investigations of zebrafish Tlr4 were performed. Using careful bioinformatics, we found an ortholog of the gene encoding Md-2 (*ly96*) in zebrafish and other ray-finned fishes. When co-transfected into mammalian cells, the zebrafish *ly96* and *tlr4ba* genes activate NF-κB signaling in response to LPS. Single-cell RNA-seq experiments on larval zebrafish revealed that the gene is expressed in a small subset of cells that express the zebrafish *tlr4*-like genes and the macrophage-specific gene *mpeg1.1* (26). This contrasts with amniotes, in which *TLR4* and *LY96* are both broadly expressed (27). Further, unlike mammalian *TLR4* and *LY96* mutants that exhibit increased resistance to systemic LPS challenge (4, 28), zebrafish larvae with loss of function *ly96* mutations are not protected from LPS toxicity. Finally, we revisited the history of the *tlr4* gene in zebrafish, finding that formation of an LPS-sensitive Tlr4/Md-2 complex is likely an ancestral feature shared by mammals and zebrafish, rather than a *de novo* invention on the tetrapod lineage. We hypothesize that zebrafish preserve an ancestral, low-sensitivity Tlr4/Md-2 complex that plays an LPS-sensing role in a small population of innate immune cells.

## MATERIALS & METHODS

### Phylogenetic reconstruction analysis

We constructed curated databases of Md-1, Md-2, Tlr4, and Cd180 protein sequences from across the vertebrates. Cd180 and Md-1 are paralogs of Tlr4 and Md-2, respectively (29). We obtained amino acid sequences of these proteins from NCBI, Ensembl, Fish1TK, amphibian transcriptomes (30–33), UniProt, and ZFIN. We constructed a multiple sequence alignment for Tlr4 and Cd180 and for Md-2 and Md-1 using MSAPROBS (34), followed by manual editing in MEGA (35). We trimmed the alignment to remove highly variable (and therefore unalignable) regions. We used PHYML (36, 37) with subtree pruning and re-grafting to construct the ML phylogeny. Pilot analyses revealed that the JTT substitution model with 8 rate categories and a floating gamma distribution parameter yielded the highest likelihood trees (38–40). An Akaike information criterion (AIC) test was used to control for overfitting (41). We rooted our trees at the duplication of these proteins in early vertebrates. Alignment figures in supplement were made with JalView (42).

### Synteny analysis

For the *ly96* synteny analysis, we used the Ensembl synteny module (43) to map homologs onto the chromosomes of species of interest. For the tlr4 synteny analysis, we took the 22 genes flanking human *TLR4 (*11 on each side) and the 22 genes flanking zebrafish *tlr4*. We used tblastn with default parameters to BLAST these sequences against 11 vertebrate genomes. We discarded all hits with e-value < 0.001 and then calculated a running average of the log (e-value) along each chromosome with a sliding window of 10,000 bases. Finally, we divided this running average by the maximum observed log (e-value)/bp value across all genomes. This value occurs for the window centered on the zebrafish *tlr4* gene. On the final relative scale, 0.0 indicates no hits observed in a given window and 1.0 is the maximum e-value per base pair. The complete analysis pipeline is implemented in a collection of shell scripts and jupyter notebooks (https://github.com/harmslab/vertebrate-tlr4-synteny/).

### Gene expression analysis

Whole 6 days post fertilization (dpf) zebrafish were euthanized by tricaine methane sulfonate overdose, flash frozen in 1 mL of Trizol (Ambion), thawed, and homogenized. Chloroform (200 μL) was added to each tube followed by mixing, centrifugation at 12,000 g for 10 minutes at 4°C, transfer of the aqueous phase to a separate tube, addition of 200 μL ethanol, and binding of sample to an RNeasy mini kit column (Qiagen). RNA was washed and eluted according to the manufacturer’s instructions and treated with RQ1 DNase (Promega). RNA was reverse transcribed into cDNA using Superscript II Reverse Transcriptase (Invitrogen) and an oligo dT (20) primer, then amplified by PCR using gene-specific primers for zebrafish *ly96* (5’ - TGTATGGCATCTGAGAAAGCAGA - 3’ and 5’ - AAGAGCAGGGGGAAACAGTC - 3’) and the housekeeping gene *b2m* (5’ - ACGCTGCAGGTATATTCATC - 3’ and 5’ - TCTCCATTGAACTGCTGAAG - 3’). PCR products were separated by electrophoresis on a 6% *bis-*Acrylamide (19:1) gel that was stained with 1X SYBR green 1 nucleic acid gel stain (Invitrogen) and imaged using an AlphaImagerHP (Alpha Innotech). The identity of the *ly96* RT-PCR product was verified by Sanger sequencing.

### Single-cell RNA-Seq

Single-cell analysis of transcription patterns of *ly96*, *tlr4ba*, *tlr4bb*, and *tlr4a1* was performed using the recently released Zebrafish Single-Cell Transcriptome Atlas (44). Briefly, dissociated cells were run on a 10X Chromium platform using v2 chemistry. Dissociated samples for each time point (1, 2 and 5 dpf) were submitted in duplicate to determine technical reproducibility. The resulting cDNA libraries were sequenced on either an Illumina Hi-seq or an Illumina Next-seq. The resulting sequencing data were analyzed using the 10X Cellranger pipeline, version 2.2.0 (45) and the Seurat software package for R, v3.4.4 (46, 47) using standard quality control, normalization, and analysis steps. We aligned reads to the zebrafish genome, GRCz11_93, and counted expression of protein coding reads. The resulting matrices were read into Seurat where we performed PCA and UMAP analysis on the resulting dataset with 178 dimensions and a resolution of 13.0, which produced 220 clusters and one singleton. Differential gene expression analysis was performed using the FindAllMarkers function in Seurat and Wilcoxon rank sum test.

### Cell Culture and Transfection Conditions

Mammalian expression vectors containing human *TLR4* and mouse *Tlr4* were obtained from Addgene (#13085 and #13086), originally deposited by Ruslan Medzhitov. Human *CD14* and *ELAM-Luc* were also obtained from Addgene (#13645 and #13029) originally deposited by Doug Golenbock. Human *MD-2* was obtained from the DNASU Repository (HsCD00439889) and contains a C-terminal V5-tag. Mouse *Md-2* (UniProt #Q9JHF9) and *Cd14* (UniProt #P10810), as well as opossum *Md-2* (UniProt #F6QBE6), *Cd14* (NCBI Accession #XP_007473804.1) and chicken *Md-2* (UniProt #A0A1D5NZX9), and *Cd14* (UniProt #B0BL87) were designed to be free of restriction sites, codon-optimized for human expression, and purchased as mammalian expression vector constructs in pcDNA3.1 (+) from Genscript (New Jersey, USA). Zebrafish *tlr4ba* and *ly96* were also obtained from Genscript in pcDNA3.1 (+). Zebrafish *tlr4bb* was a gift from Carol Kim. We re-cloned this protein from its original vector into pcDNA3.1 (+) to limit variability in expression due to differences in vector size and promoter.

Human embryonic kidney cells (HEK293T/17, ATCC CRL-11268) were maintained up to 30 passages in DMEM supplemented with 10% FBS at 37° C with 5% CO_2_. For each transfection, a confluent 100 mm plate of HEK293T/17 cells was treated at room temperature with 0.25% Trypsin-EDTA in HBSS and resuspended with an addition of DMEM + 10% FBS. This was diluted 4-fold into fresh medium and 135 µL aliquots of resuspended cells were transferred to a 96-well cell culture treated plate. Transfection mixes were made with 10 ng of *tlr4*, 1 ng of *cd14*, 10 ng of *ly96*, 1 ng of *Renilla*, 20 ng of *ELAM-Luc*, and 58 ng of pcDNA3.1 (+) per well for a total of 100 ng of DNA, diluted in OptiMEM to a volume of 10 µL/well. To the DNA mix, 0.5 µL per well of PLUS reagent was added followed by a brief vortex and room temperature incubation for 10 min. Lipofectamine was diluted 0.5 µL into 9.5 µL OptiMEM per well. This was added to the DNA + PLUS mix, vortexed briefly and incubated at RT for 15 min. The transfection mix was diluted to 65 µL/well in OptiMEM and aliquoted onto a plate. Cells were incubated with transfection mix overnight (20-22 hrs) and then treated with LPS. *E. coli* K-12 lipopolysaccharide (LPS) (tlrl-eklps, Invivogen) was dissolved at 5 mg/mL in endotoxin-free water, and aliquots were stored at −20°C. To avoid freeze-thaw cycles, working stocks of LPS were prepared at 10 μg/mL and stored at 4°C. To prepare treatments, LPS was diluted in 25% phosphate buffered saline and 75% DMEM. Cells were incubated with treatments for 4 hr. The Dual-Glo Luciferase Assay System (Promega) was used to assay Firefly and Renilla luciferase activity of individual wells. Each NF-κB induction value shown represents the Firefly luciferase activity/Renilla luciferase activity, normalized to the buffer treated transfection control to compare fold-change in NF-κB activation for treatments.

### Generation of mutant zebrafish

Zebrafish experiments were approved by the University of Oregon Institutional Animal Care and Use Committee. Chop Chop (http://chopchop.cbu.uib.no) was used to design a guide RNA (gRNA) targeting the first exon of zebrafish *ly96* (si:dkey-82k12.13, GRCz11). A gRNA template was generated by a template-free Phusion polymerase (New England Biolabs) PCR reaction using a scaffold primer (5’-GATCCGCACCGACTCGGTGCCACTTTTTCAAGTTGATAACGGACTAGCCTTATTTTAACTTGCTATTTCTAGCTCTAAAAC-3’) and an *ly96*-specific primer (5’-AATTAATACGACTCACTATAGGGTATCAGATATGGCGCTTGTTTTAGAGCTAGAAATAGC-3’), then cleaned using the QIAquick PCR Purification Kit (Qiagen), transcribed using a MEGAscript kit (Ambion), and purified by phenol-chloroform extraction and isopropanol precipitation. Cas9 RNA was made by linearizing the pT3TS-nls-zCas9-nls plasmid (41) with XbaI, purifying it using the QIAquick Gel Extraction Kit (Qiagen), performing an in vitro transcription reaction using the T3 mMESSAGE kit (Invitrogen), and purifying the RNA using the RNeasy Mini kit (Qiagen). AB strain zebrafish embryos were microinjected at the one cell stage with 1-2 nL of a mixture containing 100 ng/µL Cas9 mRNA, 50 ng/µL gRNA, and phenol red, and raised to adulthood. Fin DNA was amplified by PCR using primers specific to the targeted region (5’-CAAATTGGATTCACAACAGAGC-3’ and 5’ - CCATGGAAAATCAATGAAAAGC - 3’). Mosaic mutants were identified based on loss of an HaeII restriction site and were outcrossed to wildtype AB zebrafish to generate heterozygotes. Fish with loss-of-function mutations were identified by Sanger sequencing and further crossed to generate three independent homozygous *ly96* mutant lines.

### Fish LPS survival assay

WT and homozygous *ly96* mutant zebrafish embryos were grown under standard conditions in separate 10 cm petri dishes at a density of one fish per mL of embryo medium (EM), with fifty fish total per dish. At 5 dpf, lipopolysaccharides (LPS) purified from *Escherichia coli 0111:B4* (Sigma L2630) was dissolved in EM and added to dishes at a final concentration of 0.6 mg/mL, and control fish were mock treated with EM alone. Dead larvae, as determined by lack of heartbeat, were counted and removed at regular intervals from 16 to 48 hours or from 16 to 72 hours after addition of LPS, at which time the experiment was terminated and surviving fish were humanely euthanized.

## RESULTS

### Zebrafish have a gene encoding MD-2

The strongest evidence against Tlr4/Md-2 performing LPS-sensing in zebrafish is the lack of Md-2. Md-2 is essential for LPS recognition by amniote Tlr4, as it contains the LPS binding pocket (Fig 1). We therefore asked whether we could find a gene encoding Md-2 in bony fishes.

We started by using the human MD-2 protein sequence to BLAST against the zebrafish genome and transcriptomes. This returned no significant hits, so we took a more phylogenetically informed strategy. Relative to humans, the earliest branching, functionally characterized Tlr4/Md-2 complex is from chicken (*Gallus gallus*). We therefore “walked out” from amniotes towards fishes, starting with amphibians. We BLASTed the human MD-2 protein sequence against the *Xenopus laevis* genome. This revealed a hit to a hypothetical protein with 30% identity (OCT74818.1). When reverse-BLASTed against the human proteome, this hit returned Md-2. To validate the amino acid sequence, we compared it to the sequences of functionally characterized Md-2 proteins from amniotes. We found that the *X. laevis* gene appeared to be N-terminally truncated. Using XenBase, we identified the full-length transcript in the transcriptome for *X. laevis*. By BLASTing against available amphibian transcriptomes (30–33), we further identified putative Md-2 proteins in *Rhinella marina, Hynobius retardatus, Odorrana margaretae*, and *Ichthyophis bannanicus (*Fig S1).

With these putative amphibian Md-2 sequences in hand, we returned to our search for a zebrafish Md-2. A BLAST against a zebrafish transcriptome using the *X. laevis* sequence revealed a likely transcript (si:dkey-82k12.13, 23% identity). We then searched additional fish transcriptomes available from the Fish-T1K project (49) and identified a set of transcripts from three species that matched Md-2 (Fig S1). The genes we identified in bony fishes that encode putative Md-2 proteins were highly diverged. On average, they exhibited only 26% identity against human Md-2, and only ∼40% identity relative to one another.

We next set out to assign the orthology of these putative Md-2 sequences. Our primary concern was that these newly identified sequences were paralogs of Md-2. We therefore built a phylogenetic tree to elucidate whether these newly identified sequences grouped with Md-2 or with its direct paralog, Md-1. We constructed an alignment of 294 Md-1 and Md-2 protein sequences sampled from amniotes, amphibians, and bony fishes and then used this to infer a maximum likelihood phylogeny (Fig 2A). The alignment is available in File S1.

**Fig 2.**
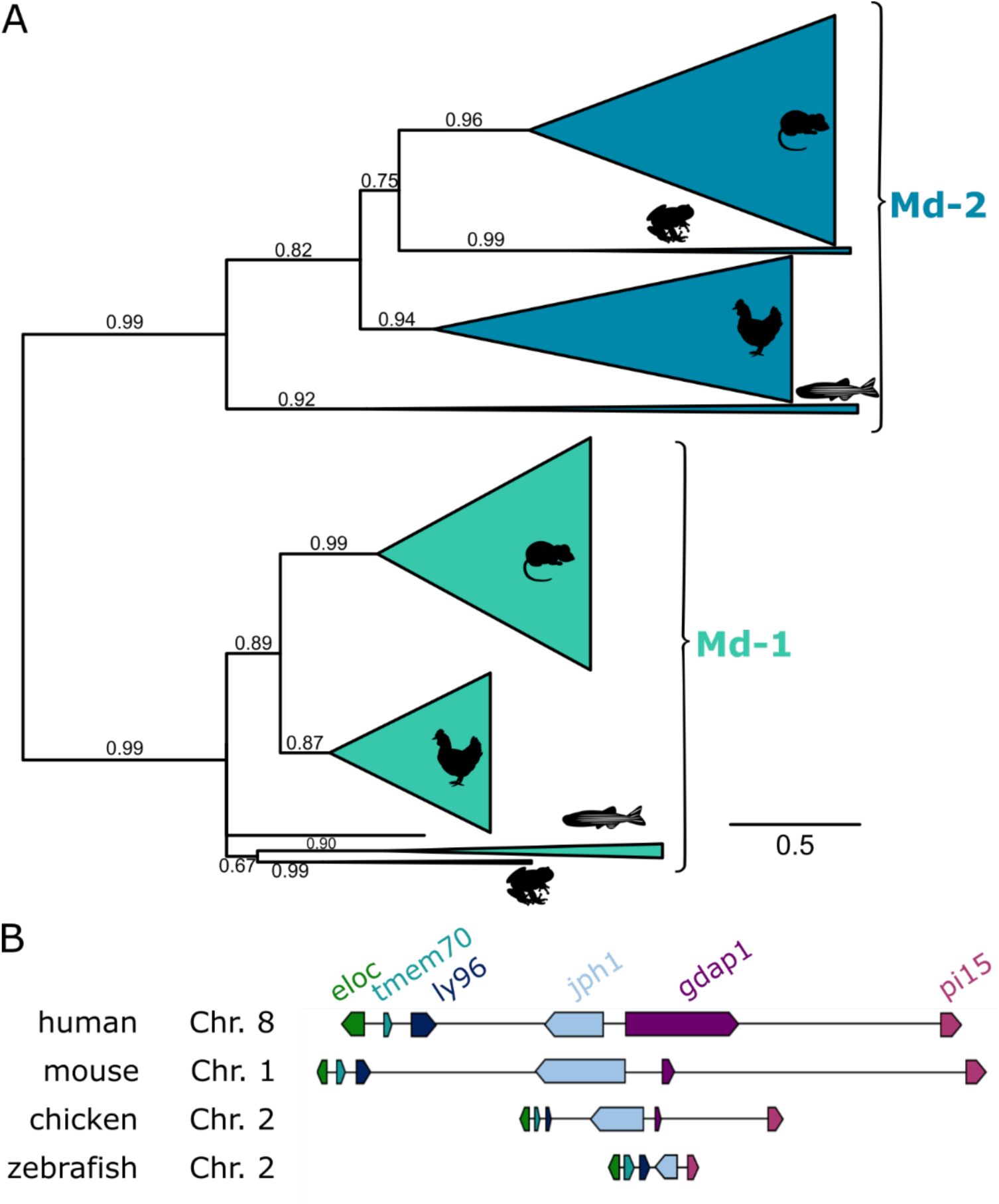
Phylogeny and synteny of the identified zebrafish protein support classifying it as an Md-2 (the *ly96* gene). A) Maximum likelihood phylogeny of Md-2 and Md-1 proteins. Wedges are collapsed clades of orthologs, with wedge height corresponding to the number of included taxa and wedge length indicating the longest branch length with the clade. Support values are SH-supports calculated using an approximate likelihood ratio test. Clades are colored to highlight Md-2 (blue) vs. Md-1 (green) classification. The taxa included in each clade are noted on the tree by silhouettes of mammals (mouse), sauropsids (chicken), amphibians (frog), and fish (zebrafish). B) Genomic organization of genes surrounding Md-2 in vertebrates. Arrows for genes represent the coding strand. Approximate distances between genes are represented by the length of line for the selected chromosome.

The putative amphibian and bony fish Md-2 sequences grouped with the tetrapod MD-2 sequences with strong support (SH = 0.99). The Md-1/Md-2 protein tree largely reproduced the species tree, with the exception of amphibians. On the Md-1 lineage, amphibians form a polytomy with fishes at the base of the tree; on the Md-2 lineage, they are placed inside the amniote clade with a relatively short internal branch. This is likely an artifact of the small number of amphibian sequences, as well as the rapid evolution of the genes along these lineages.

Overall, the tree is consistent with a single gene duplication event sometime before the evolution of bony vertebrates. Then, Md-1 and Md-2 were preserved along most descendant lineages. This said, the protein sequences of Md-1 and, particularly, Md-2 are evolving rapidly. The total branch lengths between the last common ancestor of Md-2 to its human and zebrafish descendants are 2.00 and 2.44, respectively. Put another way, the average site in the Md-2 sequence has changed its amino acid ∼2 times over the last 430 million years. Only 7 of 160 positions in MD-2 are universally conserved across the clade.

To cross-validate the orthology of this newly identified gene, we next examined its locations in the *X. laevis* and *D. rerio* genomes. We found that the synteny is consistent with other bony vertebrates (Fig 2B). In five genomes sampled from across bony vertebrates— including *X. laevis* and *D. rerio—*the gene encoding Md-2 is located between *tmem70* and *jph1b* (Table S1). This provides strong evidence that these amphibian and fish genes are, in fact, orthologous to the amniote gene encoding Md-2. By convention, the gene encoding Md-2 is known as *ly96*. We therefore refer to this gene as zebrafish *ly96* hereafter.

Due to the genome duplication event that occurred along the zebrafish lineage (50), we also looked for a second copy of *ly96*. We examined the genomic location of the *jph1a* paralog, but we were unable to identify an additional gene with any similarity to *ly96*. It appears that an inversion may have occurred in this region, complicating identification by synteny alone. This said, no additional transcripts were identified within the zebrafish transcriptome with similarity to the identified zebrafish *ly96* sequence. This is consistent with a loss of the duplicate copy of this gene.

Finally, we attempted to identify an Md-2 sequence from earlier-diverging lineages including Chondrichthyes (cartilaginous fishes) and Agnatha (jawless fishes). Despite extensive BLASTing, we were unable to identify an Md-2 protein sequence or *ly96* gene in either lineage. This is consistent with *ly96* arising after the divergence of cartilaginous and bony fishes (∼470 million years ago), but before the divergence of bony- and ray-finned fishes (∼435 million years ago). The sequence resources for cartilaginous and jawless fishes remain relatively sparse, however, so we cannot exclude an earlier origin for *ly96*.

### Zebrafish transcribe *ly96* in innate immune cells

We next asked whether zebrafish express *ly96*. To do so, we used the recently released Zebrafish Single-Cell Transcriptome Atlas (44). This dataset consists of single-cell RNASeq transcriptomes for 44,102 individual cells extracted from 1, 2 and 5 dpf zebrafish. The gray points in Fig 3A and 3B shows the entire Atlas: each point is a cell, plotted such that cells with similar transcription profiles appear near one another. Cluster identity can be established by examining differentially expressed transcripts and using these marker genes to assess cell type expression *in vivo* (44); this provides a means to assess which cell types express *ly96* simply by asking which clusters possess *ly96* transcripts.

**Fig 3.**
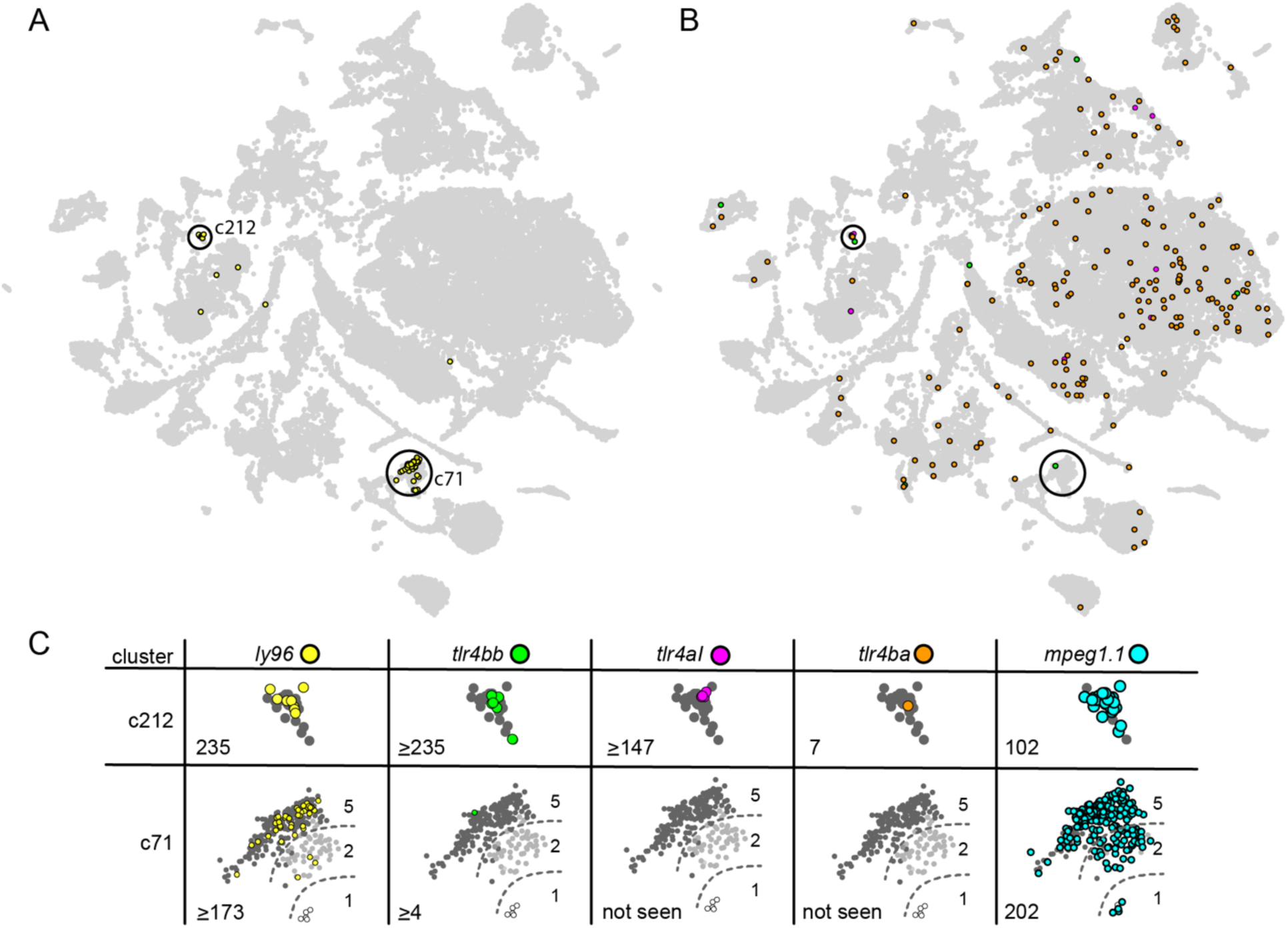
*ly96* and *tlr4* genes are expressed in macrophage cells. Each point in these plots is an individual cell characterized by single-cell RNA-Seq. The distance between the cells corresponds to the relative difference in their transcriptional profiles (44). A) Yellow points indicate cells expressing *ly96*, gray points show all 44,102 cells in the data set. The two clusters in which *ly96* is expressed (c71 and c212) are highlighted with black circles. B) Colored points indicate cells expressing *tlr4bb* (green), *tlr4al* (magenta), or *tlr4ba* (orange); gray points and circles are identical to panel A. C) Enlarged views of clusters c212 and c71, separated by gene of interest. This includes the genes shown in panels A and B, as well the macrophage marker, *mpeg1.1* (light blue) (26, 51). The number in the bottom left of each table entry is the expression level of the gene within the cluster divided by its expression level in all other cells in the dataset. If there was no expression in cells outside the cluster, expression within the cluster was divided by the detection threshold (0.001) giving a minimum estimate for the enrichment. The background cells are now colored by the developmental stage from which the cell was isolated: 1 dpf (white), 2 dpf (light gray), or 5 dpf (dark gray). The dashed lines in shown on c71 are approximate divisions between the age-dependent sub-clusters of c71.

We found that *ly96* is expressed in two clusters, denoted “c71” and “c212” (Fig 3A, yellow points). Both of these clusters are annotated in the Atlas as putative macrophage cells based on their transcription profiles (44). *ly96* is highly enriched in these clusters relative to other clusters. This can be measured by taking the ratio of the average expression level of *ly96* for the cells in the cluster relative to the average expression level of *ly96* in all other cells. This ratio is 235 for cluster c212 and ≥173 for cluster c71, indicating that *ly96* is highly enriched in these clusters. For comparison, the well-established macrophage marker *mpeg1.1* (26, 51) has ratios of 102 and 202, respectively, for these same clusters (Fig 3C).

We next investigated the expression of the *tlr4bb*, *tlr4al*, and *tlr4ba* genes. We found that *tlr4bb* and *tlr4al* had quite limited expression patterns (Fig 3B, green and pink), while *tlr4ba* was expressed broadly (Fig 3B, orange). All three *tlr4* genes were found in cluster c212, but only *tlr4bb* was found in cluster c71 (Fig 3C).

The Atlas also has the potential to reveal time-course information for the expression of these genes, as it contains cells isolated from fish at 1, 2 and 5 dpf. We therefore shaded the cells within clusters c71 and c212 by their developmental time point (Fig 3C). Cluster c212, where we observed overlapping expression for *ly96* and all three *tlr4* genes, consists entirely of cells isolated from 5 dpf zebrafish (Fig 3C). Cluster c71 has three discrete sub-clusters corresponding to the age of the fish from which the cell was extracted. We see no *ly96* in the 1 dpf sub-cluster, a small amount in the 2 dpf sub-cluster, and the highest level in the 5 dpf sub-cluster (Fig 3C). Likewise, *tlr4bb* is expressed in the 5 dpf sub-cluster but no others. For comparison, the macrophage marker *mpeg1.1* is found in all cells within c71 and c212, regardless of the age of the fish from which the cell was extracted.

These observations suggest that *ly96* and all three *tlr4* genes are expressed together in a subset of macrophage cells by 5 dpf (Fig 3C, c212). Samples of later time points would be necessary to establish if these genes are at their full expression level by 5 dpf, or if their expression level and cell-type specificity continues to change as the fish develop.

### Zebrafish Tlr4a/Md-2 can activate NF-κB in response to lipopolysaccharide

Given the low sequence similarity between the zebrafish Md-2 protein and its amniote orthologs, it was not clear that the zebrafish Md-2 would be capable of mediating the Tlr4 response to Md-2. We therefore turned to an *ex vivo* cell culture assay to assess the ability of the zebrafish Md-2 to partner with zebrafish Tlr4a and Tlr4b for LPS activation. In this assay, we co-transfected genes encoding complex components into HEK293T cells and then used luciferase to quantify NF-κB output in response to exogenously applied LPS (6).

We started by co-transfecting cells with zebrafish *ly96* and zebrafish *tlr4ba* or *tlr4bb* and then measuring NF-κB activation in response to exogenously applied LPS. We saw no activation (Fig 4A). This result was unsurprising, as this experiment attempted to activate a Tlr4/Md-2 complex without Cd14—an important peripheral protein that brings LPS to Tlr4/Md-2 complexes in amniotes, dramatically increasing the NF-κB response (Fig 1) (52–56). We thus co-transfected *tlr4ba* or *tlr4bb* with zebrafish *ly96* and human *cd14*. In this context, we observed potent activation of NF-κB in response to LPS for *tlr4ba*, but not *tlr4bb (*Fig 4A). To verify that the activation of Tlr4a required Md-2, rather than merely Cd14, we tested the activation of Tlr4 and Cd14 without transfecting *ly96*—this complex did not respond to LPS (Fig 4A). We then verified that the zebrafish Tlr4a/Md-2 complex, complemented with human Cd14, exhibited a dose-dependent response to LPS (Fig 4B). The concentration of LPS needed for activation of the zebrafish Tlr4a/Md-2 complex was much higher than that needed for activation of the human proteins in these cells, but consistent with what has been observed for other species (57).

**Fig 4.**
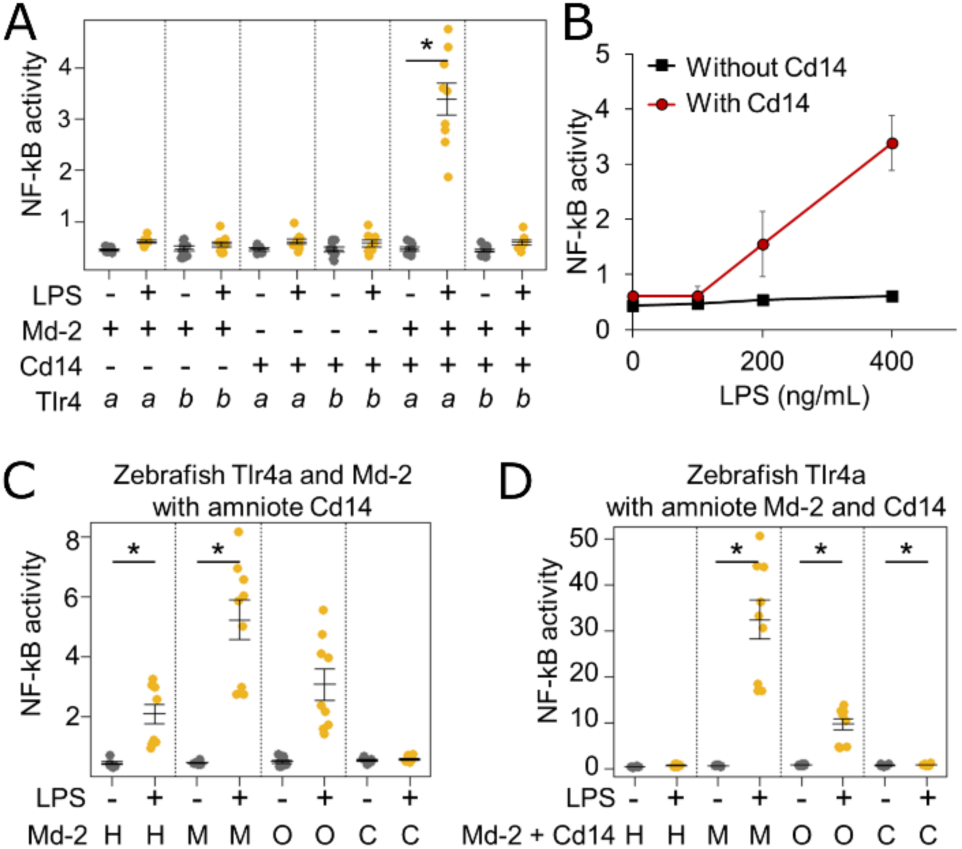
LPS activates the zebrafish Tlr4a/Md-2 in a functional assay. A) Activation of zebrafish Tlr4a and Tlr4b in the presence and absence of zebrafish Md-2 and human CD14. Points are the technical replicates from three biological replicates. Bold lines are the mean of the biological replicates. Error bars are a standard error on the mean of the biological replicates. B) Dose-dependence of LPS response by zebrafish Tlr4a/Md-2 in the presence (red) and absence (black) of human CD14. C) Zebrafish Tlr4a/Md-2 complemented with Cd14 proteins from amniotes. D) Zebrafish Tlr4a complemented with species-matched Md-2/Cd14 pairs taken from amniotes. Statistically significant differences (single-tailed Student’s t-test) are noted on each panel (* p < 0.05)

Our results support the hypothesis that zebrafish Tlr4a/Md-2 can activate in response to LPS; however, this could only be done with the presence of a supporting mammalian protein (human Cd14). To determine if this was an artifact of the human protein, we tested the LPS activation of Tlr4a/Md-2 in the presence of human, mouse, opossum, and chicken Cd14. We found that all but the chicken Cd14 were able to support the activation of the complex (Fig 4C). Thus, the activity of the zebrafish Tlr4a/Md-2 complex does not depend exclusively on human Cd14 but can instead be supported by diverse Cd14 molecules. Given the importance of Cd14 in this assay, we looked for evidence of a zebrafish *cd14* gene; however, we were unable to locate such a gene. The inability to detect a *cd14* in fish may be due to rapid evolution of this gene since the most recent common ancestor, or, alternatively, Cd14 may have arisen as a supporting molecule for LPS-recognition after the divergence of tetrapods. The requirement for Cd14 in these experiments could be a problem with the heterologous cell line (these experiments were done in human cells) or a missing alternate secondary cofactor (such as a fish LPS binding protein).

Finally, to see if zebrafish Tlr4a behaved similarly to amniote Tlr4, we investigated whether Md-2 from other species could act in concert with zebrafish Tlr4a. We co-transfected *tlr4a* with human, mouse, or opossum *ly96* genes. We saw complementation by both mouse and opossum Md-2 for LPS activation of zebrafish Tlr4a (Fig 4D). This suggests that the requirements for activation by the Tlr4/Md-2 complex have been conserved for over 400 million years and are shared across bony vertebrates.

### Md-2 is not required for LPS-induced death in 5 dpf larval zebrafish

We next probed the physiological role of Md-2 in LPS-induced septic shock in larval 5 dpf zebrafish. We first treated 5 dpf larval WT zebrafish with LPS and followed their survival over time. No treated WT fish survived more than 48 hours; the median survival time was 30 hrs (Fig 5A). As a control, we also tested the LPS response for *myd88^-/-^* fish. As has been observed previously (22), these showed a modest but significant increase in survival (Fig 5A). This was consistent with LPS inducing a response that depends in part on a *myd88* dependent pathway.

**Fig 5.**
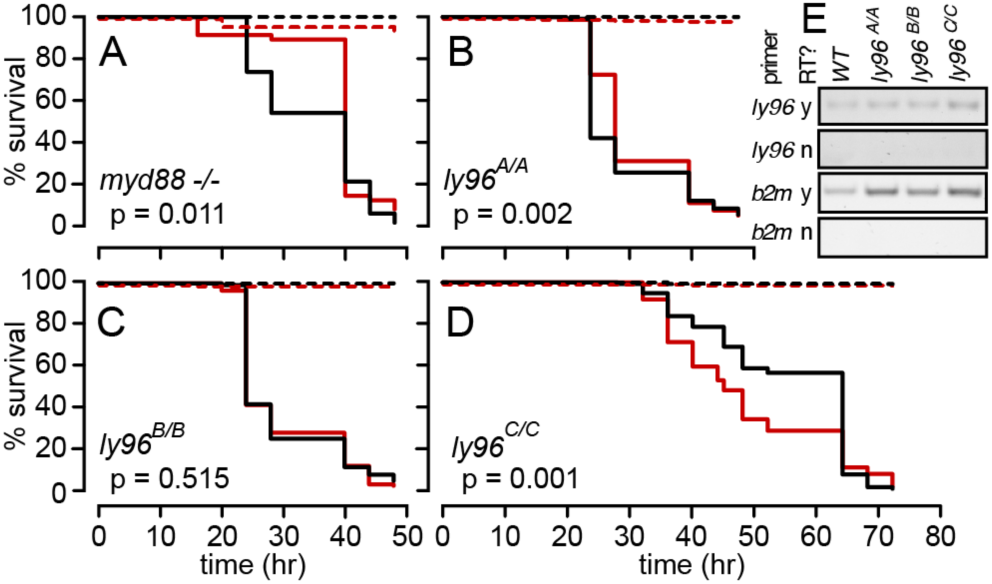
*ly96* mutations only moderately affect LPS survival in larval zebrafish. A-D) Curves show survival of wildtype (black) and mutant (red) zebrafish in the presence of 0.6 mg/mL LPS (solid line) or mock treatment (dashed line). The genotype is indicated on each panel. The p-value was determined by comparing the matched survival curves by a log-rank Mantel-Cox test. The experiments shown in panels A-C were performed in parallel, while the experiments in panel D were performed at a later date with an LPS lot that showed lower potency, necessitating a longer treatment time. Panels A-D represent averages of one, five, five, and three experimental repeats, respectively. Panel E shows mRNA transcript level for each mutant zebrafish. Rows use different primers (*ly96* or *b2m*) with and without reverse transcriptase (RT). Columns show fish genotype.

To test the role of Md-2 in this response, we used CRISPR-Cas9-based mutagenesis to establish three independent zebrafish lines with mutations in the first exon of the *ly96* gene. The mutations were expected to induce a loss of function through removal of the start codon (*ly96^A/A^*) or through a frame shift and premature stop codon (*ly96^B/B^* and *ly96^C/C^*) (Table S2). Using RT-PCR primers downstream of the targeted region, we demonstrated that *ly96* mRNA is expressed in mutant larval zebrafish (Fig 5E).

We then tested the three *ly96* mutant zebrafish lines for their susceptibility to LPS-induced septic shock. The results were mixed. Compared to matched WT controls, *ly96^A/A^* zebrafish survived for slightly longer (Fig 5B); *ly96^B/B^* zebrafish survived similarly (Fig 5C), and *ly96^C/C^* zebrafish survived shorter (Fig 5D). This is consistent with some other pathway rather than Tlr4/Md-2 being the primary route for LPS-induced death in larval zebrafish.

### The zebrafish *tlr4* paralog arose after the evolution of *ly96*

Finally, we revisited the idea that the evolutionary history of zebrafish *tlr4* genes implies that they do not act as LPS-sensing molecules. Previous authors suggested that an ancestral *TLR* gene duplicated in the ancestor of bony vertebrates (∼450 million years ago), and that the two paralogs were then differentially lost on the mammalian and bony fish lineages (21), respectively. This early divergence, before the evolution of *ly96*, may suggest very different functional roles for mammalian versus fish *tlr4*s.

We set out to better resolve when the zebrafish *tlr4* paralog arose relative to its mammalian counterparts, particularly with regard to the evolution of *ly96*. As with our analysis of Md-2, we started with a phylogenetic tree and then turned to synteny. For the phylogenetic tree, we constructed a multiple sequence alignment containing 263 Tlr4 sequences and 190 Cd180 protein sequences as an outgroup. (Cd180 is the most closely related paralog to Tlr4) (58). The alignment is available in File S2. In the resulting maximum likelihood tree, Tlr4 and Cd180 form distinct, well-supported clades (Fig 6A). Within the Tlr4 clade, zebrafish Tlr4a and Tlr4b are part of a monophyletic group with other Tlr4s.

**Fig 6.**
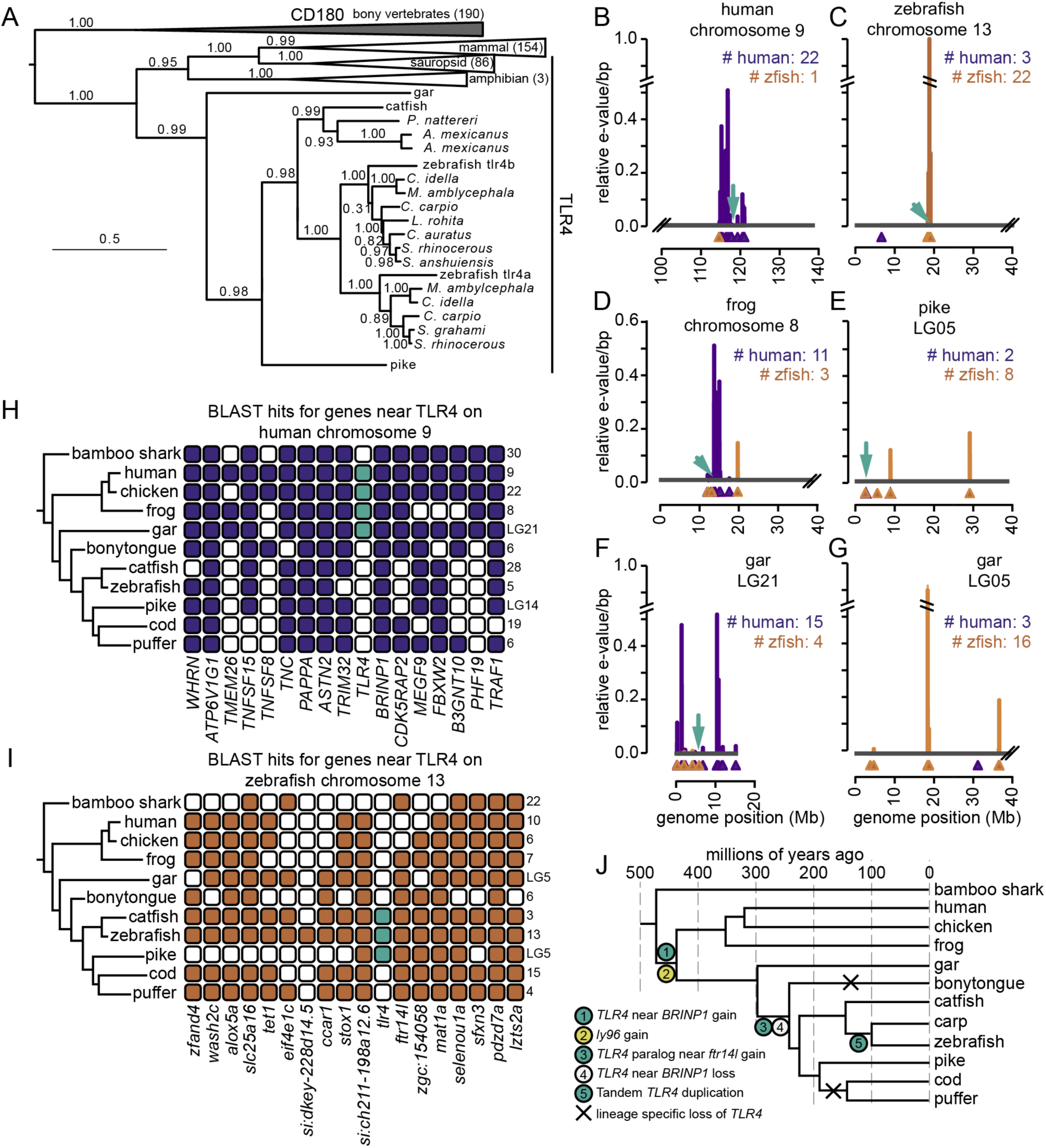
Zebrafish *tlr4* paralogs evolved within the ray-finned fishes. A) Maximum likelihood phylogeny for 453 Tlr4 and Cd180 protein sequences. SH supports are indicated on the tree. Wedges are clades, with the length indicating the maximum branch length from the ancestor of the clade. The taxonomic distribution and number of genes within each wedge are indicated on the plot. B-G) Hits for human (purple) and zebrafish (orange) gene sets on six representative chromosomes taken from five species. The species and chromosome are indicated at the top of each plot. The x-axis denotes position on the chromosome. Triangles indicate gene start positions. The green arrow indicates the location of a Tlr4 gene. The y-axis is a running average of the BLAST e-value for each gene set along the genome (see methods). The numbers on the plot indicate the number of human and zebrafish hits within the region shown. H,I) Each row shows the chromosome with the most BLAST hits from the human (panel H) or zebrafish (panel I) gene set. Columns indicate specific genes from the set, with names denoted below. A colored square indicates a gene found somewhere on the chromosome. A green square is a *tlr4* gene. The species tree is shown on the left; the chromosome number is on the right. J) Schematic representation of a plausible scenario for the history of the *tlr4* gene. Times are taken from Hughes et al. (59) and timetree.org (60).

We next investigated the genomic context for *tlr4* genes in eleven genomes, each with a complete chromosome assembly (Table S3). We selected a set of 22 genes flanking human *tlr4* and a set of 22 genes flanking the three zebrafish *tlr4* genes (Table S4). Notably, there were no shared homologs between the sets, demonstrating the radical difference between the genomic contexts of human and zebrafish *tlr4*. We then used these sets of genes to BLAST each of the eleven genomes and calculated a running average for the BLAST e-values along each chromosome. This allowed us to assess the overall similarity of genomic regions to either the human or zebrafish *tlr4* context. Fig 6B-G shows representative traces for six chromosomes taken from five species, with results for all genomes in Fig S2-S13. We were able to distinguish two distinct contexts for *tlr4* genes. In some organisms—human and frog, for example—*tlr4* is surrounded by hits from the human gene set (Fig 6B and D). In other organisms—zebrafish and pike, for example—*tlr4* is surrounded by hits from the zebrafish gene set (Fig 6C and E).

To place our results in their evolutionary context, we plotted our BLAST output against the phylogeny for our chosen species. For each species, we displayed the chromosome with the most hits from the human set (Fig 6H) and the chromosome with the most hits from the zebrafish set (Fig 6I). We made an exception for the pike, displaying the chromosome with the *tlr4* gene (linkage group 5), not the chromosome with the most zebrafish hits (linkage group 6). We indicated whether a gene from the human or zebrafish set was seen somewhere on that chromosome by coloring the square corresponding to that gene.

Four species had *tlr4* in a human-like context: human, chicken, frog and gar. None of these species—including the gar—had a duplicate copy of *tlr4* in a zebrafish-like context. The human-like context of the gar gene is shown Fig 6F, while the lack of *tlr4* in the most zebrafish-like region of the gar genome is shown in Fig 6G. The remainder of the ray-finned fishes had *tlr4* in either a zebrafish-like context (catfish, zebrafish, and pike) or had no *tlr4* gene at all (bonytongue, cod, and puffer).

The most parsimonious history consistent with the observed distribution across genomes is shown in Fig 6J. In this scenario, *tlr4* arose in a genomic context similar to the one preserved in humans. This occurred after the divergence of bony and cartilaginous fishes (∼475 million years ago), but before the divergence of ray-finned and lobe-finned fishes (∼430 million years ago). The ancestral genomic context was preserved in tetrapods, including humans. It was also maintained in the ray-finned fishes for ∼130 million years, as indicated by the location of the *tlr4* gene in the gar genome. Then, sometime between 300 and 250 million years ago, the *tlr4* gene was both duplicated into the genomic context observed in zebrafish, as well as lost from the ancestral context. Finally, between 150 and 100 million years ago, a tandem duplication occurred within the *Cypriniformes* fishes, leading to the tandem copies of *tlr4* observed in zebrafish, carp, and other *Cypriniformes* fishes.

This revised evolutionary history places the evolution of the zebrafish *tlr4* paralogs much later than was previously hypothesized (21). Importantly, the duplication of TLR4 occurred *after* the evolution of Md-2, meaning that the formation of the Tlr4/Md-2 complex likely pre-dates the duplication event. Thus, the interaction with Md-2 and the ability to activate with LPS were an ancestral feature of zebrafish Tlr4 rather than something that could only be gained in parallel along the tetrapod and bony fish lineages.

## DISCUSSION

Our observations led us to reevaluate the decade-old idea that Tlr4 does not participate in the LPS-induced inflammatory response in zebrafish. We have identified the zebrafish gene encoding the Tlr4 co-receptor Md-2 (*ly96*). The gene, like *tlr4ba* and *tlr4bb,* is transcribed in zebrafish cells that transcribe a collection of macrophage genes. In concert with zebrafish Tlr4a, zebrafish Md-2 is capable of activating NF-κB signaling in an *ex vivo* functional assay. Finally, a careful phylogenetic analysis suggests that the mammalian and zebrafish *tlr4* genes are not as evolutionarily distinct as previously thought. While not direct orthologs, the zebrafish paralogs evolved well after *ly96* and likely preserve an ancestral LPS recognition activity.

Our work demonstrates that, given the correct context, zebrafish Tlr4a and Md-2 form a functional complex that recognizes LPS and activates NF-κB signaling. Further, the molecular basis for the interaction between the partners appears to have been conserved for the last 450 million years - zebrafish Tlr4a is compatible with mouse and opossum Md-2 (Fig 4D). This is despite the fact that the orthologous proteins from each species have only ∼20% identity at the amino acid sequence level. The simplest explanation for this observation is that the ability of Tlr4/Md-2 to activate in response to LPS is an ancestral feature of the protein complex that has been conserved across the bony vertebrates—from mammals to bony fishes.

We have not shown, however, that LPS-induced Tlr4/Md-2 signaling actually occurs in zebrafish. Our two attempts to do so—our cell culture functional assay and analysis of zebrafish *ly96* loss of function mutants—both gave mixed results. We will discuss each in turn.

### LPS activation of Tlr4a/Md-2 in human cells requires supporting molecules

In our functional assays, we had to add a mammalian Cd14 to activate NF-κB signaling through zebrafish Tlr4a/Md-2 (Fig 4C). In amniotes, Cd14 delivers LPS directly to Md-2 (Fig 1). We could find no ortholog to *cd14* in the zebrafish genome.

One possibility is that the human cell line used for the functional assays is missing some critical component for the delivery of LPS and assembly of the active dimer. Tlr4, Md-2, and Cd14 are the necessary and sufficient set of amniote proteins that confer an LPS-dependent NF-κB response in HEK293T cells. It could be that some other non-homologous protein plays the role of Cd14 in zebrafish.

Another possibility is that LPS is not a zebrafish Tlr4a/Md-2 agonist *in vivo*. We showed that we can activate the complex in a human cell line given an appropriate delivery molecule and a high enough LPS concentration. But, under physiological conditions, the Tlr4a/Md-2 complex could respond to some other chemically similar ligand. This would not be surprising: changes in ligand specificity have been observed across Md-2 in the amniotes (61). There is also some evidence that zebrafish Tlr4a may be antagonized by LPS *in vivo* (20). This would be compatible with another ligand activating the complex and LPS competing and activating at a lower level than can be achieved by the native ligand.

Finally, our observation that Tlr4a activates NF-κB with both mouse and opossum Md-2 directly contrasts previous work that showed the complex could not activate (Fig 4C) (20, 21). The key difference between our experiments and those done previously is the sequence of *tlr4a* used. Previous investigators used a construct that was ∼75 amino acids shorter than tetrapod Tlr4s. This construct is missing both the signal peptide required to target Tlr4a to the cell surface and a region of the protein that is likely critical for Md-2 binding (Fig S13). In contrast, we used a full-length ORF (ENSDART00000044697.6, GRCz10). The difference in our constructs arises because the previous analysis relied on cDNA that, apparently, captured an alternate splice variant of *tlr4a*.

### Multiple pathways contribute to LPS-induced death in larval zebrafish

Larval zebrafish *ly96* loss of function mutants did not exhibit appreciably altered death rates upon exposure to LPS compared to WT (Fig 5B-D). This is consistent with a previous morpholino study that knocked down *tlr4a* and observed no change in sensitivity to LPS (20). This contrasts with mice, however, where knockout of *LY96* is protective against endotoxic shock (4) and disruption or knockout of *Tlr4* leads to hypo-responsiveness to LPS (1, 62).

We cannot rule out the possibility that this lack of response is due to an experimental artifact. First, zebrafish may have retained a second copy of the *ly96* gene from the teleost genome duplication that maintained function even after deletion of the targeted copy. We were unable find any evidence of such a gene; however, the challenge of finding the original *ly96* gene means that we cannot rule this out. A second possibility is that the mutants that we generated may not represent a complete loss of function. For example, use of a potential alternative start codon 17 amino acids downstream of the normal start codon could produce a truncated protein (*ly96^A/A^*). Although this would be missing N-terminal amino acids that are known to be critical for Md-2 function in other systems (Table S2), these amino acids may not be necessary in zebrafish. Finally, we tested a single developmental time point. It could be that Tlr4a/Md-2, while expressed in larvae (Fig 3C), is not yet a large player in LPS sensing. Further experiments on zebrafish at different time points may help clarify this point.

Another challenge is that LPS-mediated death is a relatively blunt instrument to test for the activity of the Tlr4/Md-2 complex. We observed that addition of LPS dramatically increased death rate (Fig 5A), even in a *myd88^-/-^* background. This indicates that at least one other non-Toll-like pathway contributes to LPS-induced. One possibility is that this occurs by intracellular sensing of LPS via caspases and inflammasomes (63). Various studies have shown inflammasome signaling to be widespread in zebrafish larvae (64) and Il-1r to be required to prevent cell death in response to infection in multiple cells (65). Intracellular sensing may be much more important in fish than mammals: zebrafish have 385 of these putative intracellular sensors whereas humans have 22 (66).

As a result of such alternate pathways, even if the Tlr4a/Md-2 complex contributes to the LPS-induced inflammatory response, removing it might not lead to a measurable difference in death rate. Compounding this difficulty, our expression analysis revealed that zebrafish *ly96* is much more restricted in its expression than the corresponding mammalian genes (27). There may, in fact, be specific subtypes of macrophages that express *ly96* and *tlr4s*—and are defective in LPS sensing in the *ly96* mutants—but remain invisible at the level of LPS-induced death. Higher-resolution studies of LPS-induced inflammation will be required to sort this out.

### Snapshot in the evolutionary history of this complex

The presence of Md-2 in zebrafish indicates that both Tlr4 and Md-2 existed, together, in the last common ancestor of bony vertebrates. Because descendants along both the tetrapod and ray-finned fish lineages activate with LPS, the ability to respond to LPS is likely an ancestral function that has been conserved for 435 million years.

That said, these proteins have evolved significantly since this shared ancestor. Along the tetrapod lineage, a supporting collection of proteins evolved. Cd14 arose through a duplication within the Toll-like receptor family and is now an essential component of the Tlr4/Md-2 complex, delivering LPS to Md-2 in a coordinated fashion (53, 54). Tetrapods also acquired Lipid Binding Protein (LBP), improving LPS delivery (52, 67). Amniotes then further adjusted the Tlr4/Md-2 pro-inflammatory response through addition of amniote-specific Damage-Associated Molecular Pattern (DAMP) molecules such as S100A9 (68), which amplify LPS-induced inflammation (69). All the while, mutations to Md-2 changed its specificity for LPS and its chemical analogs (70). For example, humans acquired unique lipid IVa antagonism sometime after the divergence of humans and mice (71, 72).

The changes that occurred along the ray-finned fish lineages are not yet clear. Did they acquire supporting LPS delivery molecules analogous to Cd14? Has the specificity of Md-2 fluctuated in ray-finned fishes as it has along the tetrapod lineage? Further work is needed to answer these questions.

We hypothesize, however, that ray-finned fishes maintain an ancestral, low-sensitivity Tlr4/Md-2 LPS sensing complex. Fish have previously been shown to be relatively resistant to septic shock (73, 74), with high concentrations of LPS needed to activate teleost leukocytes (25, 75–77). This parallels the observation that early diverging tetrapods, such as amphibians, also require high doses of LPS to trigger an inflammatory response (78). This could be explained if ray-finned fishes do not have specialized machinery to deliver LPS to the complex, but instead use Tlr4/Md-2 as a simple LPS sensor. Other observations consistent with a relatively primitive Tlr4/Md-2 LPS response in zebrafish are the fact that Tlr4 was lost, independently, along multiple fish lineages (20, 21, 73) (Fig 6J), as well as the existence of parallel LPS sensing pathways in zebrafish (64). If the Tlr4/Md-2 complex is peripheral to the LPS response in ray-finned fishes, it could be lost with minimal fitness consequences. In contrast, Tlr4/Md-2 became progressively more central to the LPS response along the mammalian lineage—and as a result has been highly conserved.

Our work suggests that we should re-visit our understanding of LPS signaling through Tlr4/Md-2 in zebrafish. We hypothesize that zebrafish preserve an ancestral, low-sensitivity Tlr4/Md-2 complex. In contrast to mammals—in which the Tlr4/Md-2 complex is the primary LPS receptor—the zebrafish Tlr4/Md-2 complex acts in parallel with several LPS-sensitive pathways, likely playing roles in a small population of innate immune cells.

## ACKNOWLEDGMENTS

We thank Kristi Hamilton and Lila Kaye for assistance with zebrafish LPS survival assays, and Rose Sockol and the University of Oregon Aquatic Animal Care Services staff for fish husbandry. We thank Prof. Carol Kim for sharing the *tlr4bb* plasmid.

## SUPPLEMENTAL MATERIAL

Fig S1: Alignment of Md-2 proteins from amphibians and various fishes

Fig S2: Human and zebrafish gene context BLAST hits on bamboo shark chromosomes

Fig S3: Human and zebrafish gene context BLAST hits on human chromosomes

Fig S4: Human and zebrafish gene context BLAST hits on chicken chromosomes

Fig S5: Human and zebrafish gene context BLAST hits on frog chromosomes

Fig S6: Human and zebrafish gene context BLAST hits on gar chromosomes

Fig S7: Human and zebrafish gene context BLAST hits on bonytongue chromosomes

Fig S8: Human and zebrafish gene context BLAST hits on catfish chromosomes

Fig S9: Human and zebrafish gene context BLAST hits on zebrafish chromosomes

Fig S10: Human and zebrafish gene context BLAST hits on pike chromosomes

Fig S11: Human and zebrafish gene context BLAST hits on cod chromosomes

Fig S12: Human and zebrafish gene context BLAST hits on puffer chromosomes

Fig S13: Comparison of zebrafish Tlr4a sequence used in this paper versus previous work

Table S1: Gene locations for *ly96* synteny analysis

Table S2: Predicted zebrafish *ly96* mutant gene products

Table S3: Genomes used for *tlr4* synteny analysis

Table S4: Genes used to BLAST for human vs. zebrafish *Tlr4* genomic context

File S1: Spreadsheet containing accession numbers and aligned sequences for Md-1 and Md-2 sequences.

File S2: Spreadsheet containing accession numbers and aligned sequences for Tlr4 and Cd180 sequences.

