## Supplemental Figures for "Identification and characterization of zebrafish Tlr4 co-receptor Md-2"

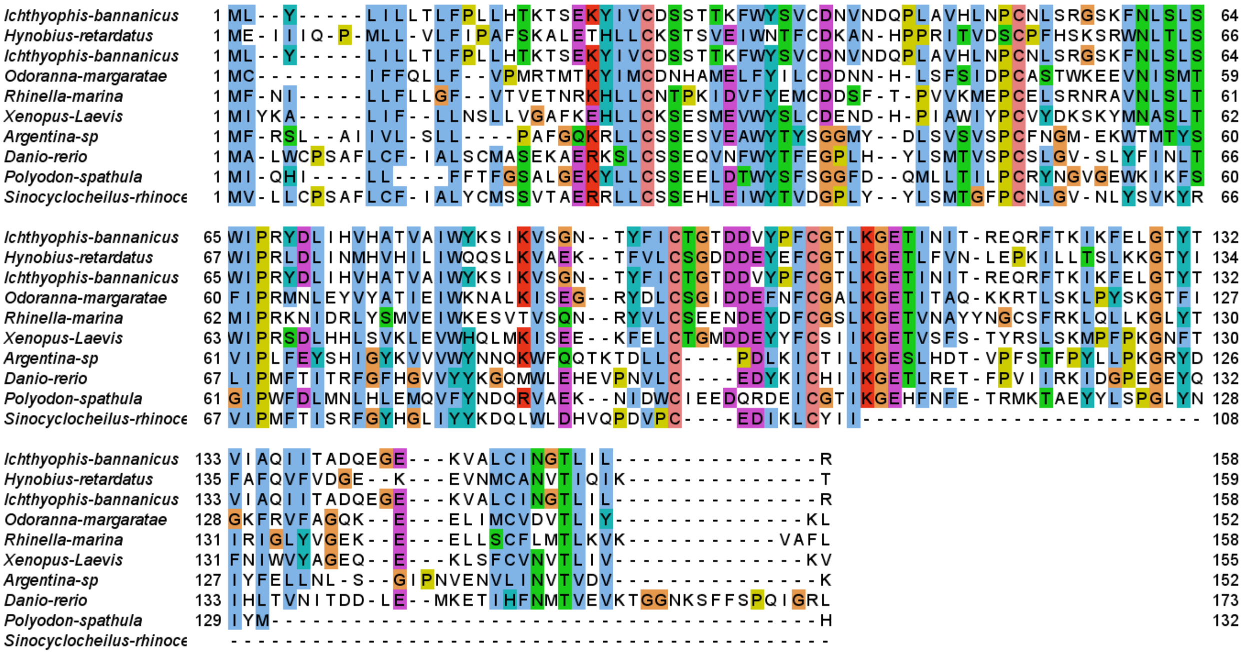


**Figure S1. Alignment of newly identified fish and amphibian Md-2 proteins**. Md-2 proteins were identified using transcriptomic data from amphibians (dryad) and fish (Fish-T1K). Sequences were aligned with MSAPROBS. Colors are used to denote chemically similar residues and conservation. Alignment was constructed with Jalview.

Fig S2: hits on bambooshark chromosomes

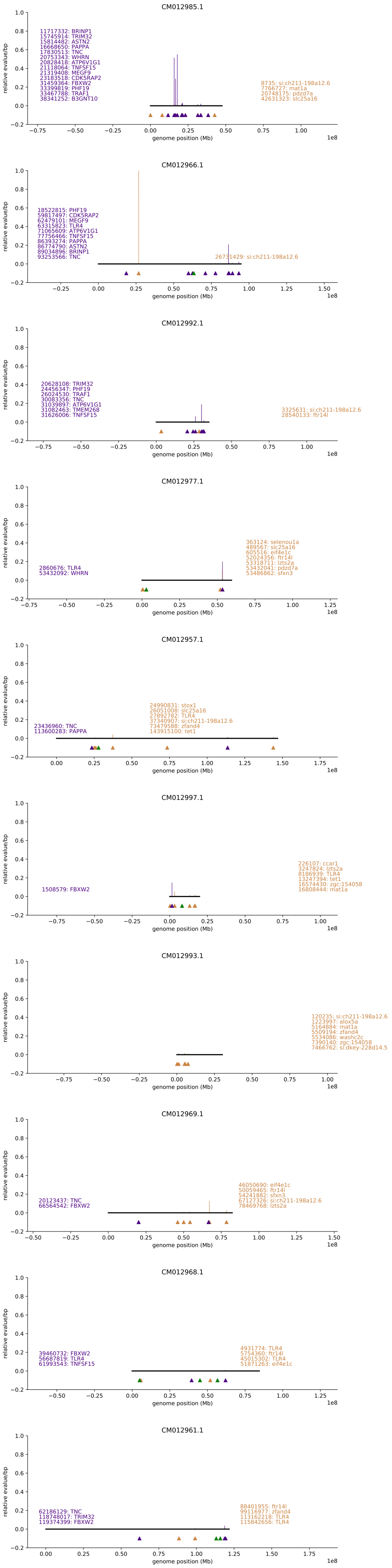

Fig S3: hits on homo chromosomes

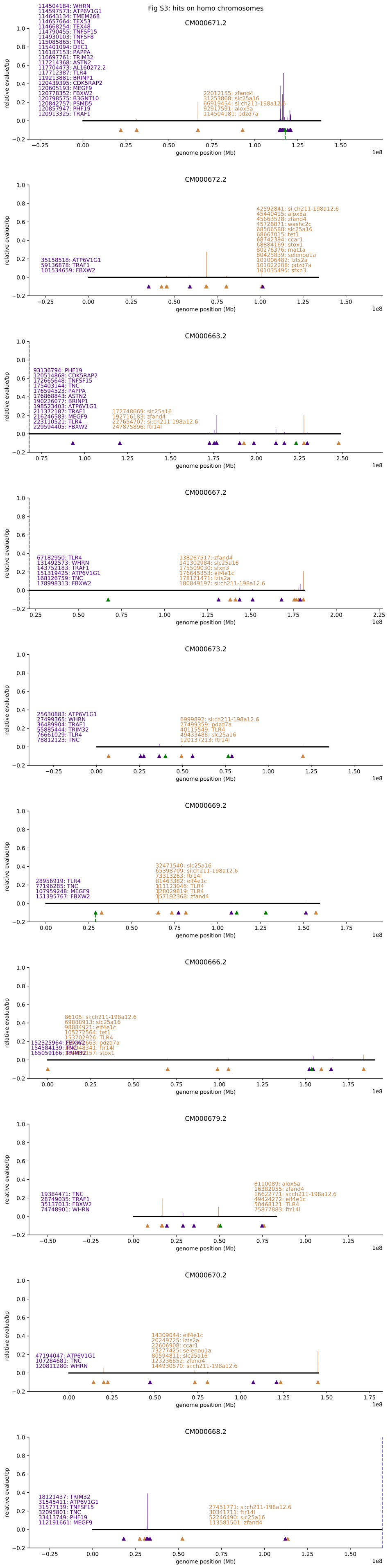

Fig S4: hits on gallus chromosomes

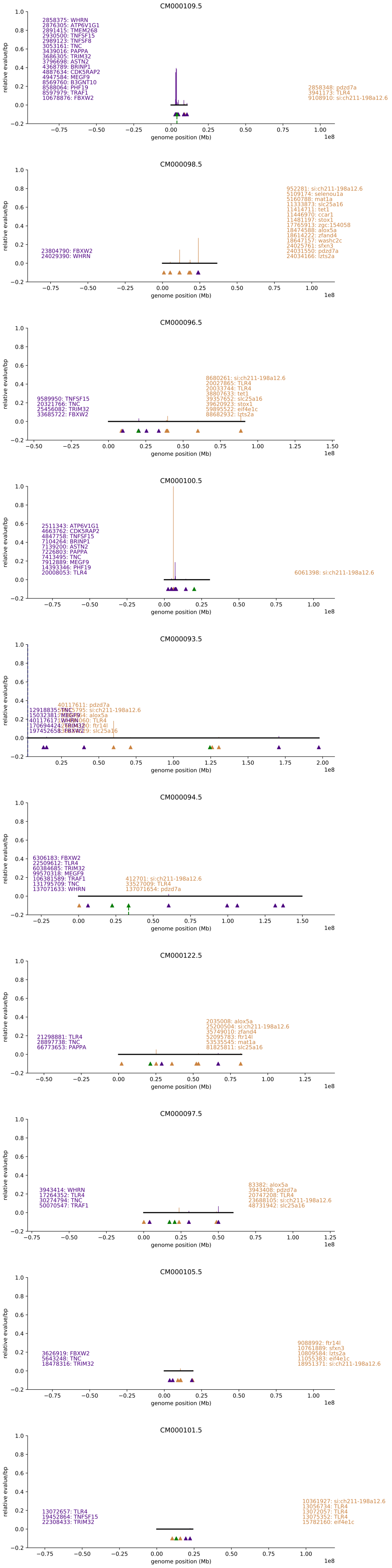

Fig S5: hits on xenopus chromosomes

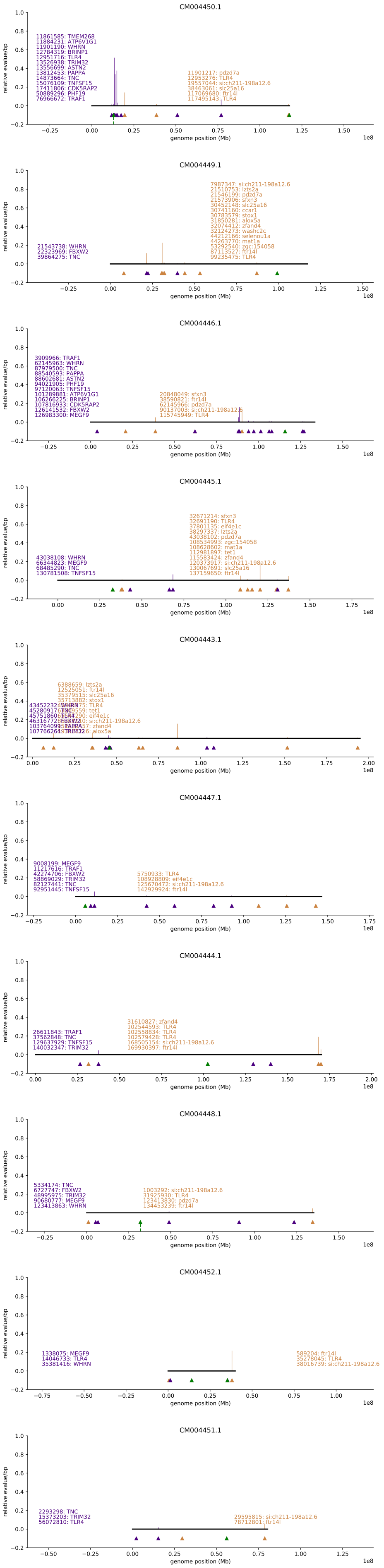

Fig S6: hits on lepisosteus chromosomes

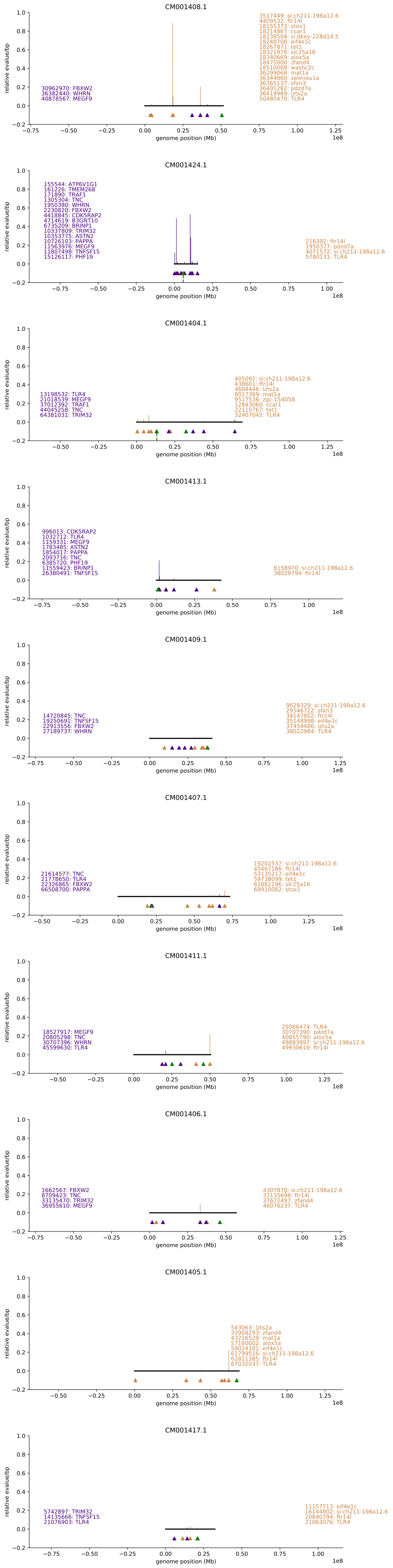

Fig S7: hits on bonytongue chromosomes

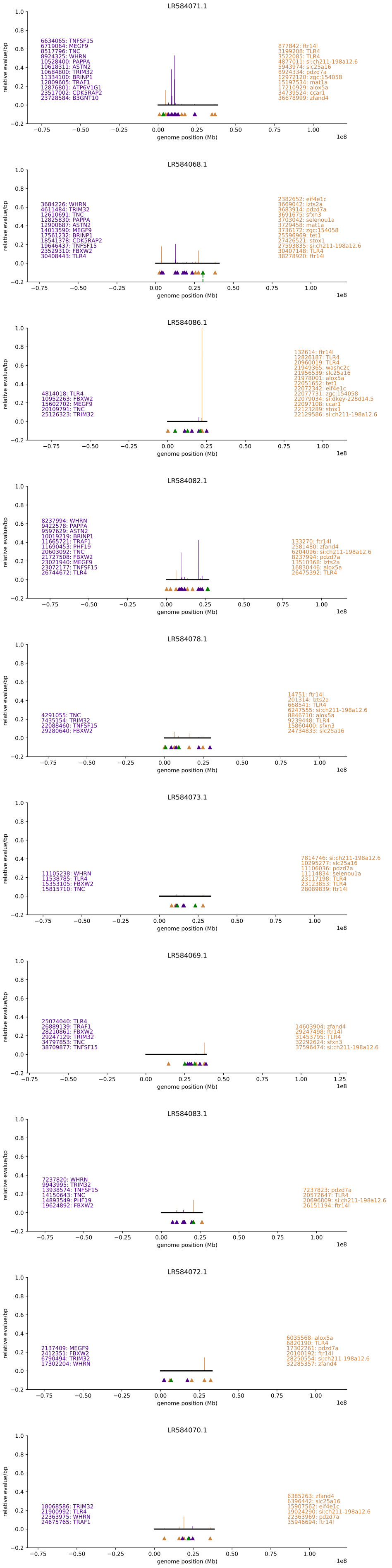

Fig S8: hits on ictalurus chromosomes

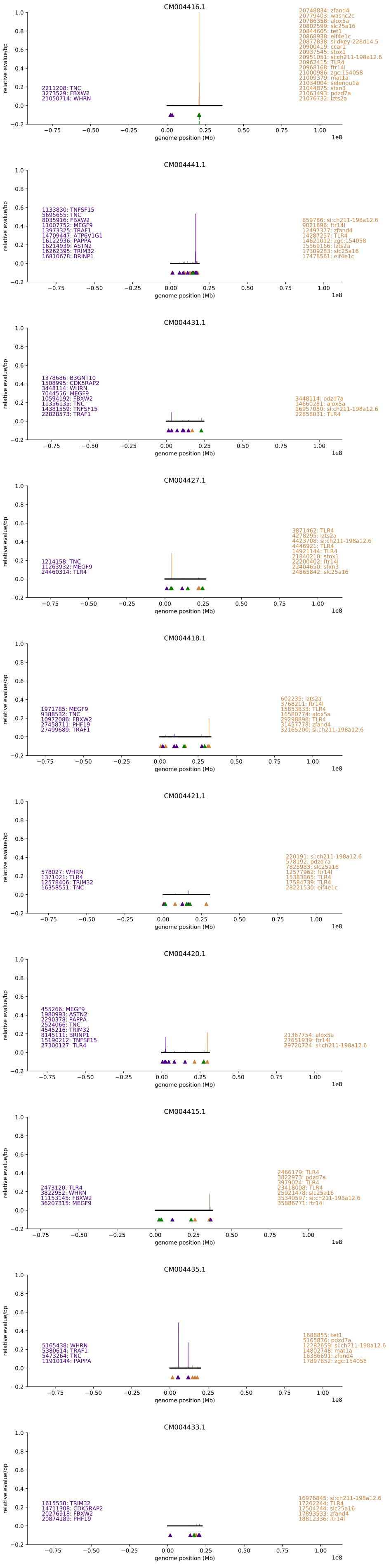

Fig S9: hits on danio chromosomes

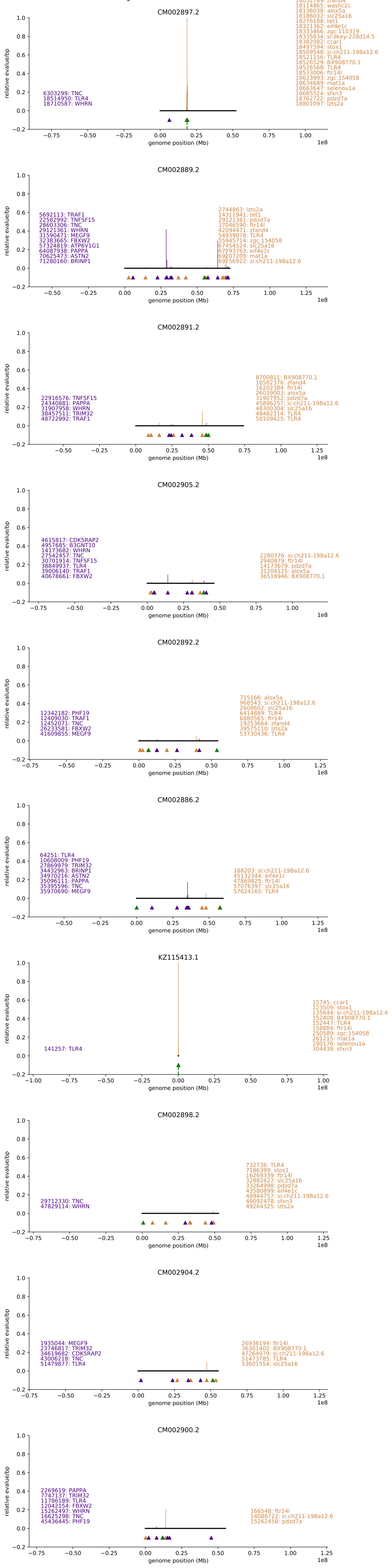

Fig S10: hits on esox chromosomes

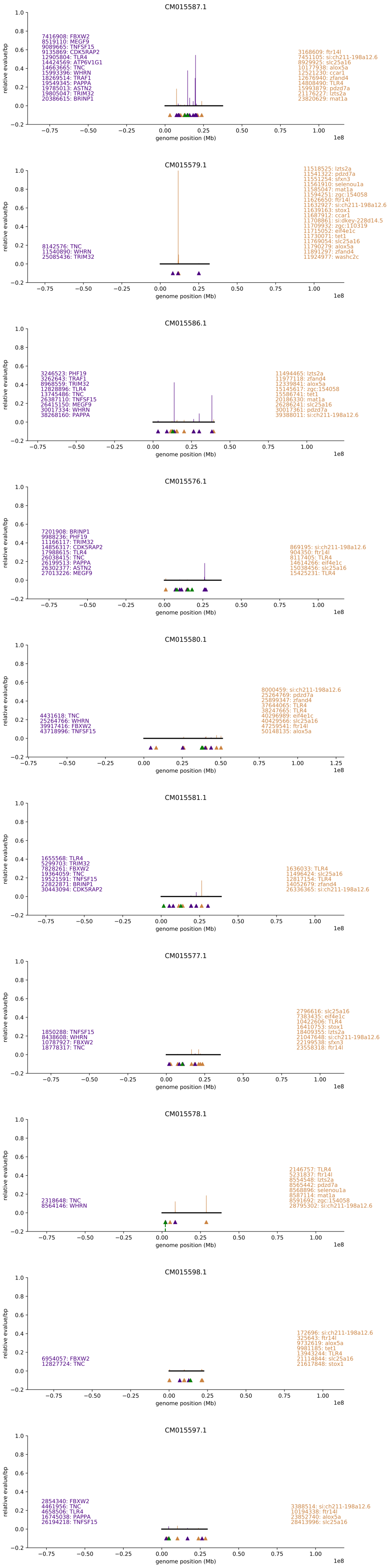

Fig S11: hits on gadus chromosomes

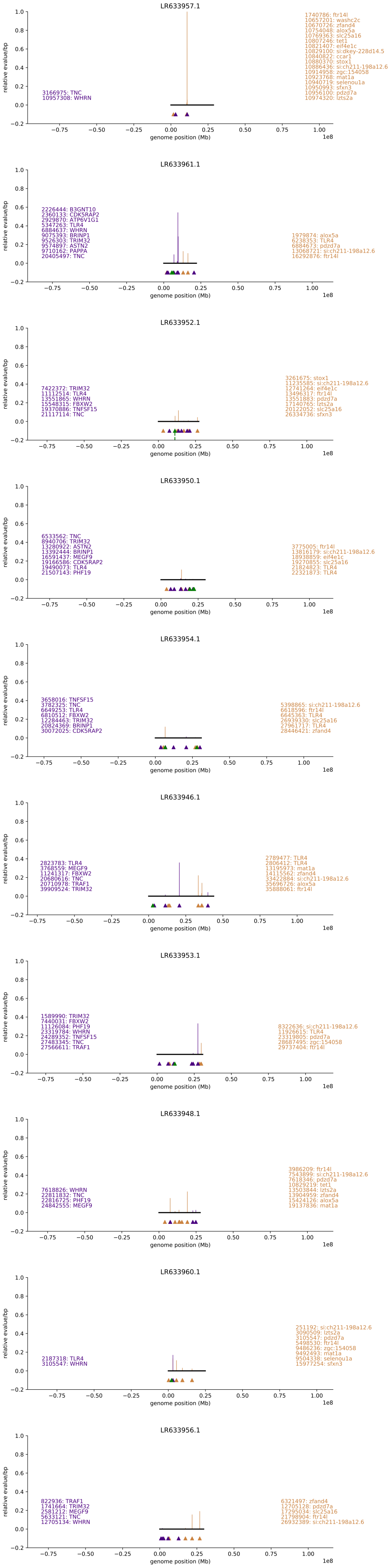

Fig S12: hits on takifugu chromosomes

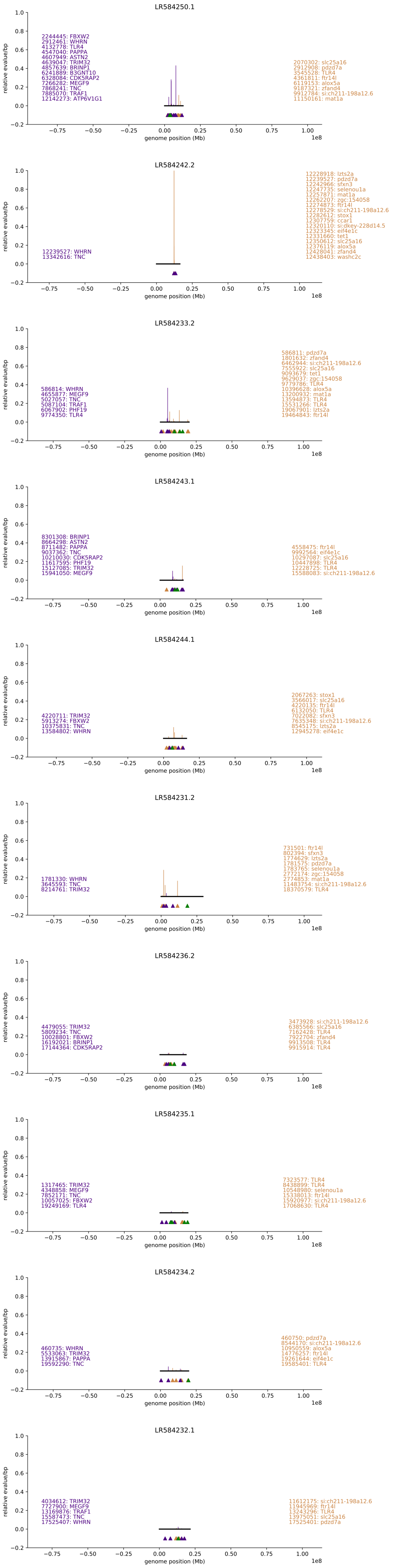

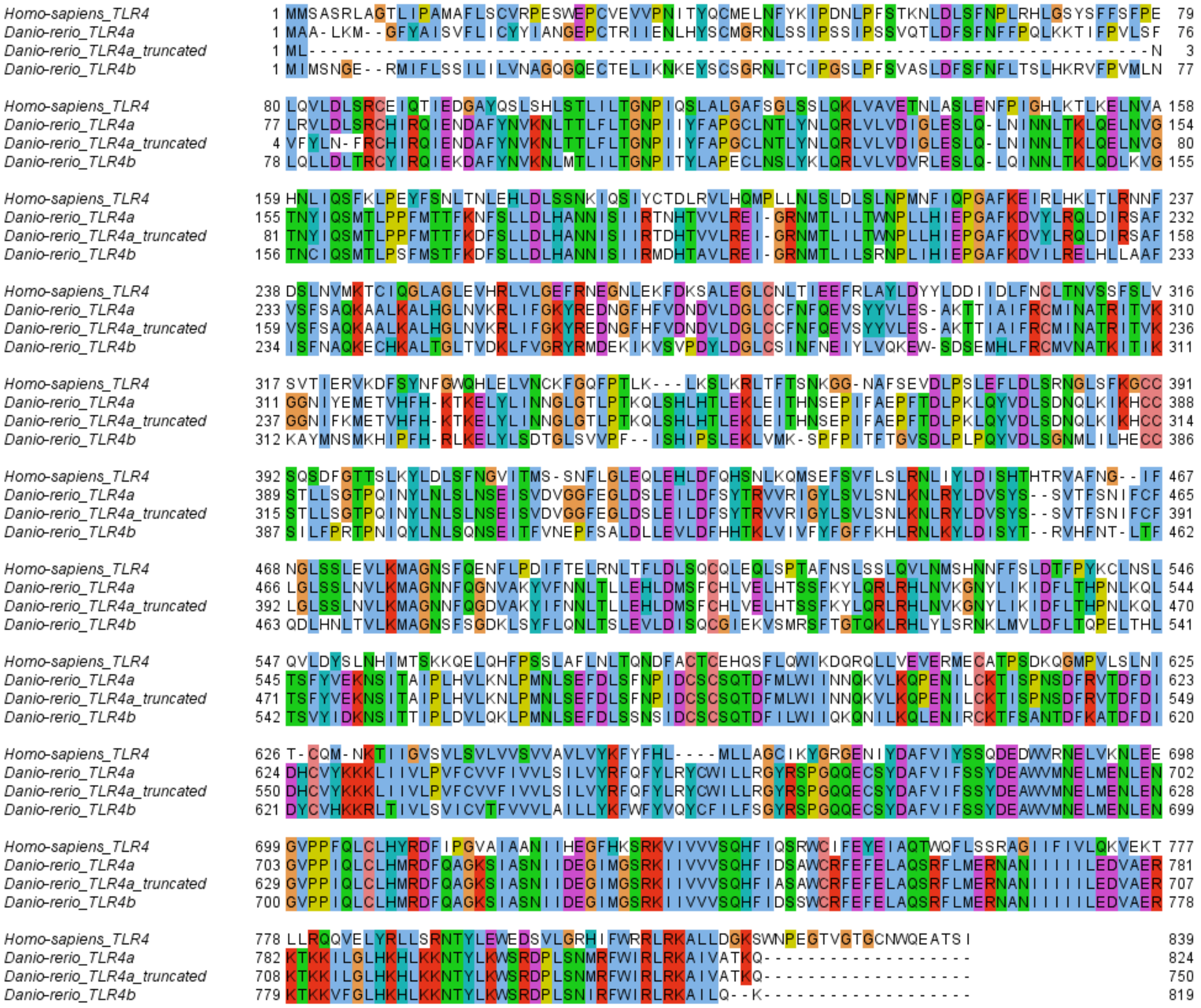


**Figure S13: Comparison of zebrafish Tlr4a sequence used in this paper versus previous work.** Sequences were aligned to human TLR4 with T-coffee. Colors are used to denote chemically similar residues and conservation. Alignment was constructed with Jalview.
